# Regulation of Oocyte Meiotic Maturation: Unraveling the Interplay between PKA Inhibition and Cdk1 Activation

**DOI:** 10.1101/2023.09.02.556023

**Authors:** Martina Santoni, Nabil Sekhsoukh, Sandrine Castella, Tran Le, Marika Miot, Enrico Maria Daldello

## Abstract

Oocyte meiosis is arrested at the first prophase stage, until hormonal stimulation triggers progression into the meiotic divisions. This process, called meiotic maturation, depends on extensive post-transcriptional events. In all vertebrates, two bottleneck events orchestrate meiosis resumption: first, the inhibition of PKA and, second, the activation of Cdk1, the master regulator of eukaryotic cell division. However, the molecular events occurring between these two steps are almost unknown. To address this issue, we took advantage of a Cdk1 inhibitor to identify the early events that depend on PKA downregulation and occur independently of Cdk1 activity. Unexpectedly, we show that accumulation of Cyclin B1 and Mos, the kinase responsible for MAPK activation in oocytes, are regulated in an opposing manner by a two-step mechanism. PKA downregulation induces first the accumulation of Cyclin B1 without any increase of its translation, independently of Cdk1 activation. Subsequently, the rate of Cyclin B1 translation increases in response to Cdk1 activation. In contrast, Mos translation begins downstream PKA inhibition, but the protein does not accumulate until Cdk1 is activated. These intertwined regulations create the positive feedback loops required for the full activation of Cdk1. Additionally, we show that two consecutive waves of translation occur during the G2-M transition, the first induced by PKA inhibition and the second by Cdk1 activation. Finally, we demonstrate that Arpp19, the only known early substrate of PKA in *Xenopus* oocytes, is not involved in the control of these early events. This study reveals that PKA downregulation promotes multiple molecular pathways that converge on the activation of Cdk1 to induce the G2/M transition in vertebrate oocytes.

## INTRODUCTION

The development of female gametes, known as oogenesis, is a crucial process for sexual reproduction. In animals, oogonia enter meiosis and arrest in prophase of the first meiotic division. During this long-lasting arrest lasting days to years, oocytes undergo an extensive phase of growth, accumulating RNAs and proteins. The full-grown oocytes then resume meiosis upon hormonal stimulation by undergoing two consecutive meiotic divisions without an intervening S-phase. Meiotic divisions are orchestrated by the activation of the Cdk1-Cyclin B complex or M-phase promoting factor (MPF), the universal cell division inducer in eukaryotes [1]. Understanding the mechanisms of MPF activation in oocytes is therefore essential to get new insight into this step fundamental for the success of fertilization and early embryogenesis. Despite its universal character, MPF is regulated in slightly different ways depending on cell type and species. This is especially true for meiotic divisions, where specific regulations of MPF enable the arrest in prophase (equivalent to a late G2-phase), then the succession of two divisions without intermediary DNA replication, and finally a new arrest, in metaphase of the second meiotic division in vertebrates. Despite the importance of these mechanisms in physiology of reproduction, how MPF activation is controlled remains unclear in oocytes.

In all vertebrate oocytes, the prophase arrest is maintained by a high level of cAMP that activates the cyclic AMP-dependent protein kinase (PKA)[2]. In *Xenopus* prophase-arrested oocytes, Cdk1 protein is present into two inactive pools: as a monomer, and as a dimer with Cyclin B2, which is inactivated by Myt1 kinase that phosphorylates Cdk1 at T14 and Y15. This stockpiled inactive complex is called pre-MPF [3]. In *Xenopus*, the release of the prophase arrest is triggered by progesterone. The signalling pathway induced by the hormone and leading to Cdk1 activation remains unclear, although it requires a decrease in PKA activity, as in all vertebrate oocytes [4]. Indeed, increase of cAMP concentration by treating oocytes with stable cAMP analogs, phosphodiesterase inhibitors, or activators of adenylyl cyclases, or by increasing PKA activity by overexpressing its catalytic subunit (PKAC) prevents Cdk1 activation and meiotic maturation induced by progesterone [5–7]. Conversely, inhibition of PKA activity by injecting either its regulatory subunit (PKAR) or its specific inhibitor (PKI) promotes meiotic maturation in the absence of progesterone [6,8]. These results demonstrate that the PKA downregulation induced by progesterone is both necessary and sufficient for Cdk1 activation in oocytes. In *Xenopus*, PKA inhibition occurs within 30 min in response to progesterone [4]. It promotes the synthesis of new proteins essential for Cdk1 activation and the meiotic maturation process, since inhibitors of protein synthesis block both progesterone- and PKI-induced meiotic maturation [9]. However, the identity of the newly synthesized proteins needed for Cdk1 activation is largely unknown. Rodents are the only known vertebrate species in which Cdk1 activation does not depend on the translation of new proteins [10]. Indeed, in mouse, the global activation of protein translation is detected only after the nuclear envelope breakdown (NEBD), the usual marker of Cdk1 activation [11].

In *Xenopus* oocytes, the full activation of Cdk1 that induces NEBD occurs 3 to 7 hours after progesterone exposure, into two steps: first a small starter amount of active Cdk1 is formed; second this Cdk1 starter activity fires the conversion of pre-MPF (Cdk1-Cyclin B2) into active MPF. This second step relies on a complex network of feedback loops involving many well-characterized kinases and phosphatases. The core of this so-called MPF autoamplification loop is the activation of Cdc25 phosphatase that dephosphorylates and activates the stockpiled pre-MPF, as well as the inactivation of Myt1, preventing the inhibitory phosphorylation of Cdk1 [12]. Once Cdk1 is fully activated, the oocyte undergoes NEBD, then enters in metaphase I (MI), achieves an asymmetric division by extruding the first polar body, enters in metaphase II (MII) and arrests again with a stable MII spindle, waiting for fertilization. While the network of Cdk1 regulators involved in the autoamplification loop is well characterized, the events required for the formation of the starting amount of active Cdk1 are still unclear. Two mechanisms have been proposed for the formation of this initial Cdk1 starter [13]. According to the first one, newly synthesized Cyclin B1 that accumulates in response to progesterone and binds to monomeric Cdk1, would form a small pool of active MPF complexes by escaping Myt1 inhibition [3,14]. According to the second one, the *de novo* translation of specific proteins would interfere with the network of kinases and phosphatases that maintains the pre-MPF inhibited, resulting in the conversion of a small amount of pre-MPF into active MPF. In the first case, the Cdk1 starter amount is formed of Cdk1-Cyclin B1 while in the second case, it consists of Cdk1-Cyclin B2. The second mechanism has been proposed to account for the action of Mos, a protein kinase that is not expressed in prophase oocytes and whose translation is induced by progesterone [15–17]. Mos activates the mitogen-activated protein kinase (MAPK) pathway (Mos/MEK/MAPK/p90^Rsk^) that would modulate the core regulators of Cdk1 [18]. Importantly, antisense oligonucleotides targeting the B-type Cyclins mRNAs or morpholino blocking Mos translation do not block MPF activation in response to progesterone, although they strongly delay the process [13].

To unravel the molecular connection between the initial drop in PKA activity, the stimulation of translation and the accumulation of new proteins, and the final activation of Cdk1, PKA substrates must be identified. We identified the first protein, Arpp19, whose phosphorylation by PKA is sufficient to block the G2/M transition [19] and whose dephosphorylation by PP2A-B55δ is required to allow the oocyte entry into M-phase [20]. Arpp19 is dephosphorylated at S109 within 30 min after progesterone exposure or PKI injection [19]. However, the events controlled by the dephosphorylation of Arpp19 that unlock a signaling pathway leading to Cdk1 activation, remain an open question. Interestingly, Arpp19 plays a key role into the second step of Cdk1 activation, the autoamplification loop. Phosphorylated at S67 by the kinase Greatwall (Gwl), it inactivates PP2A-B55δ, that opposes Cdk1-Cyclin B2 activation [21–24].

Despite the discovery of this important PKA substrate, several critical and currently unanswered questions need to be addressed to understand the molecular cascades initiated by PKA inhibition and leading to the formation of an active MPF primer. Are there any PKA substrates other than Arpp19? Which new proteins are accumulated in response to PKA inhibition? Is their accumulation regulated by translation and/or protein degradation? What is the connection between these novel proteins and PKA substrates (upstream), and Cdk1 (downstream)? Is the starting Cdk1 activity generated by Cyclin B1 or by CyclinB2? Experimental approaches to answer these questions are not trivial. Indeed, it is mandatory to unambiguously distinguish the molecular events triggered by PKA inhibition, which occur upstream and independently of first step of Cdk1 activation.

In this work, we took advantage of the Cdk1 inhibitor, Cip1, to rigorously define the molecular events triggered by PKA downregulation that occur independently of Cdk1 activation (“early events”). Having identified three of these events, we evaluated the involvement of Arpp19 in these pathways. Our work highlights that multiple and parallel molecular pathways are triggered by PKA downregulation to control Cdk1 activation and the G2/M transition in vertebrate oocytes. It reveals the unexpected regulation of the accumulation of Cyclin B1 and Mos depending on an interplay between translation and stabilisation, and the existence of additional PKA effectors in addition to Arpp19 to control the meiotic G2/M transition.

## RESULTS

### A temporal classification of the events during the meiotic G2/M transition

Meiotic resumption is initiated by PKA downregulation that occurs within 1 hr after hormonal stimulation and is followed several hours later by the activation of Cdk1. The molecular pathways connecting PKA inactivation to Cdk1 activation involve extensive post-transcriptional rearrangements, which are characterized by accumulation/degradation and phosphorylation/dephosphorylation of key proteins. To understand the regulation of these pathways, it is required to identify the molecular events that occur downstream of PKA inhibition but independently of Cdk1 activation, that we call the “early events”, and discriminate them from the “late events” that require Cdk1 activation. These two types of events are usually classified according to their temporal occurrence, before or after NEBD, triggered by Cdk1 and generally considered to be contemporary with its activation. NEBD is characterized by the appearance of a white spot at the animal pole of the oocyte that is the only visible morphological marker of the G2/M transition in *Xenopus* oocyte. Hence, it is the marker generally considered as a good dividing line separating what is dependent on Cdk1, occurring from NEBD onwards, from what is independent of it, occurring before NEBD. Fig. 1A recapitulates the main molecular events taking place in response to progesterone. Cdk1 activity was measured by the phosphorylation of endogenous PP1 at T320, a specific substrate of Cdk1 used for *in vitro* Cdk1 kinase assays, and MAPK activity by the phosphorylation of Erk1/2 [25–27]. We detected several events occurring before the pigment re-arrangement that marks NEBD: the begin of Gwl phosphorylation, Cyclin B1 and Mos accumulation, the cytoplasmic polyadenylation element (CPE)- binding protein 1 (CPEB1) destabilization, Cdk1 and MAPK activation. Before NEBD, Cdk1 activity is clearly detectable although it is still phosphorylated at Y15 (Fig. 1A). Therefore, this initial Cdk1 activity does not result from the dephosphorylation of the inactive Cdk1-Cyclin B2 complexes stockpiled in the oocyte. It results from the association of free Cdk1 molecules with newly synthesized Cyclin B1. At NEBD, the dephosphorylation of Cdk1 on Y15 takes place marking the full activation of Cdk1, which is marked by a further increase in PP1 phosphorylation at T320. Interestingly, the events that begun before NEBD continue and intensify at NEBD, as the accumulation of Cyclin B1 and Mos, the activation of MAPK and the degradation of CPEB1. Finally, other events occur after NEBD, as the accumulation of Cdc6 or the degradation and then re-accumulation of Cyclin B1 and B2 (Fig. 1A).

**Fig. 1.**
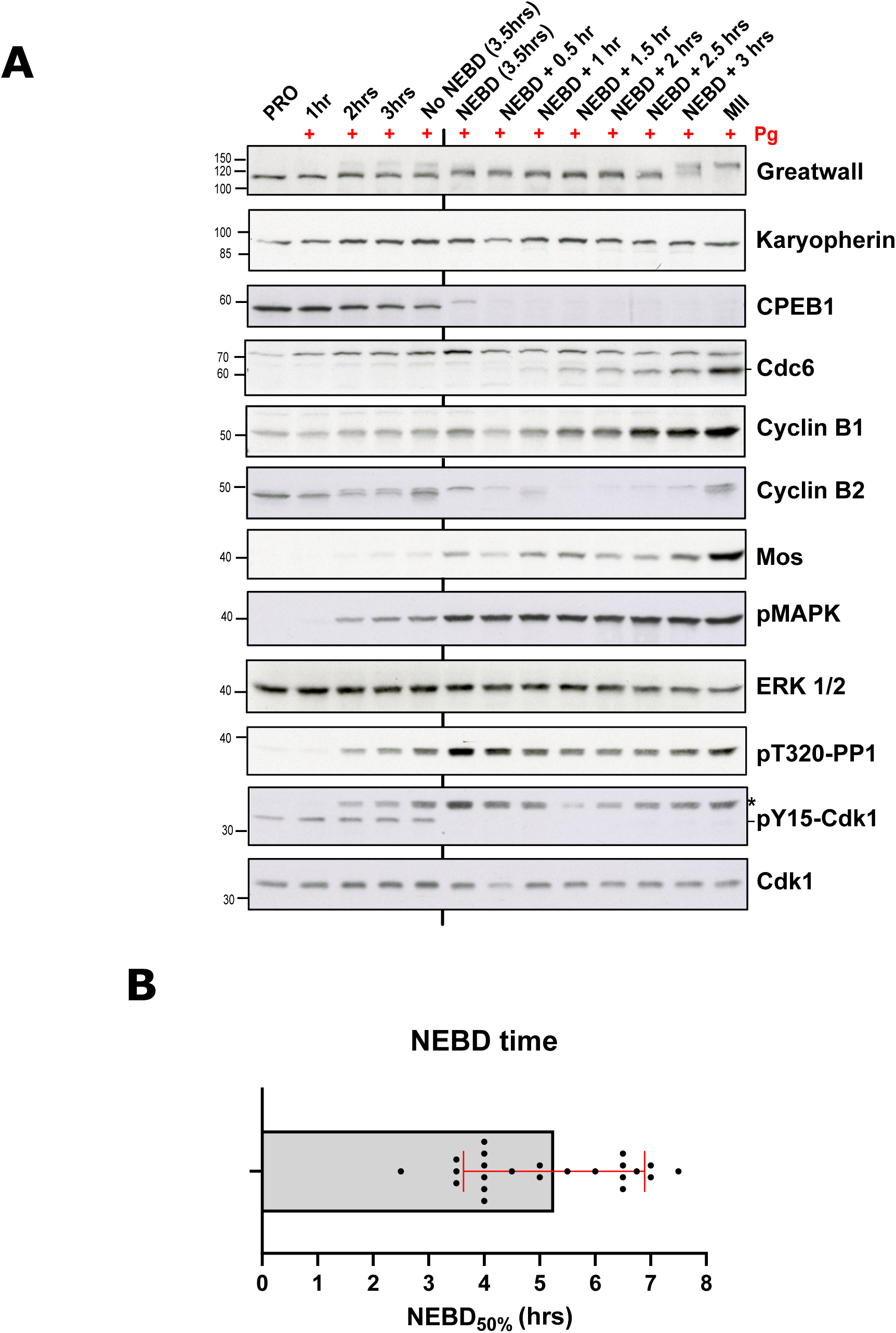
Western blot analysis of key players of meiotic maturation. A. Prophase oocytes (PRO) were incubated with progesterone (Pg) to induce meiotic maturation. 5 oocytes were collected at different times after progesterone addition. At 3.5 hrs, 50% of the oocytes underwent NEBD (NEBD50%). At NEBD50%, 5 oocytes that underwent NEBD (NEBD) or not (no NEBD) were collected. A vertical black line marks the sample collected after NEBD. A re-synchronization was done at NEBD: oocytes undergoing NEBD within a time-window of 15 min were grouped together and incubated for the indicated times before being collected. After lysis, oocyte extracts were analyzed by western blot using antibodies directed against the proteins indicated on the right. Molecular weights (kDa) are indicated on the left. *: Band representing the signal from the previously used anti-pT320-PP1 antibody. B. The time of NEBD50% of the oocytes from the 24 *Xenopus* females used in this study. Each dot represents a female.

These results clearly demonstrate that Cdk1 is active before the appearance of the white spot at NEBD, illustrating the limitations of an approach entirely based on a temporal classification of the oocytes following progesterone addition and using the NEBD white spot as a marker. Indeed, Cdk1 activity is detectable 1.5 hr before the time of NEBD 50% (time at which 50% of oocytes have completed NEBD), in oocytes without any pigment rearrangement. Therefore, during this 1.5-hour time window between the first activation of Cdk1 and its full activation at NEBD, it is impossible to distinguish which events are upstream or downstream the first activation of Cdk1. Moreover, the time of NEBD varies among oocytes obtained from the same female, and pools of oocytes from different females have different distributions of NEBD time. Indeed, among the females used for this research, the average time of NEBD 50% was 5.3 hrs ± 1.6 hrs (Fig. 1B). Consequently, another experimental approach is needed to discriminate which events occurring upstream of NEBD are dependent or not on Cdk1 activity.

### A biochemical classification of events during meiosis based on the inhibition of Cdk1

To identify the events induced by PKA inactivation independently of Cdk1 during meiosis, a biochemical strategy based on the injection of the Cdk1 inhibitor, Cip1, was implemented (Fig. 2)[14]. We cloned the *Xenopus* homolog of Cip1, produced a GST-Cip1 fusion protein and purified it. It is hereafter referred to as “Cip1”. The injection of Cip1 fully suppressed Cdk1 activation and the G2/M transition (Fig. 2). Indeed, oocytes did not undergo NEBD and Cdk1 remained fully phosphorylated at Y15 (Fig. 2A-B). The absence of Cdk1 activity in Cip1-injected oocytes was confirmed by an *in vitro* kinase assay (Fig. 2C). Furthermore, the formation of the starting amount of Cdk1 activity was also prevented by Cip1 injection, as demonstrated by the absence of phosphorylation of PP1 at T320 (Fig. 2B-C). Hence, the injection of Cip1 allows to reveal events induced by progesterone independently of Cdk1 activity by fully inhibiting its activation.

**Fig. 2.**
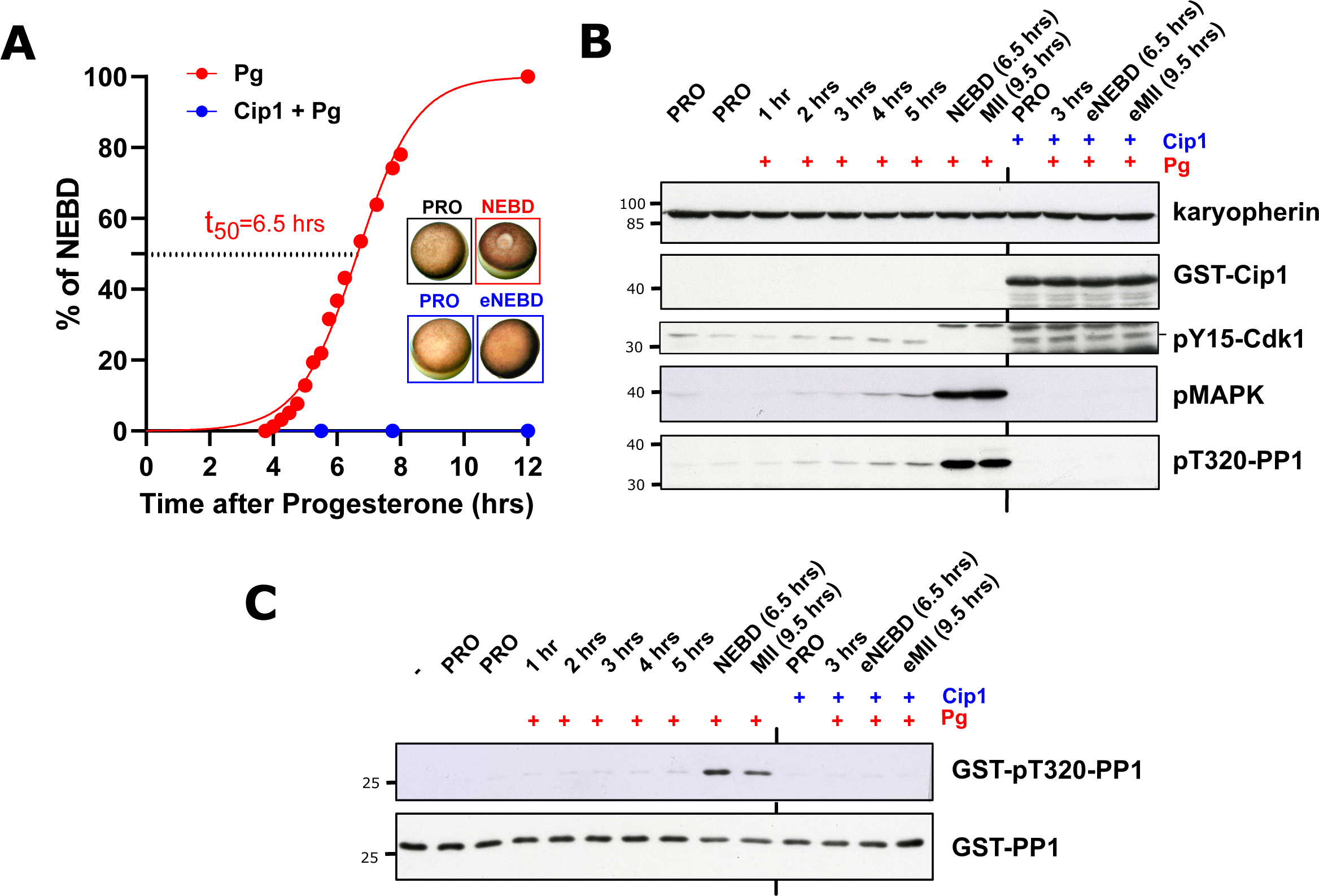
Cip1 injection blocks meiotic maturation. A. Oocytes were injected (Cip1+Pg – blue line) or not (Pg – red line) with Cip1. After overnight incubation, oocytes were incubated with progesterone (Pg) to induce meiotic maturation. The cumulative time of NEBD is shown as well as some representative pictures of oocytes collected at the prophase stage (PRO) or at NEBD. Oocytes injected with Cip1 were collected at the time when 50% of the non-injected oocytes underwent NEBD (equivalent NEBD – eNEBD). B. Western blot analysis of the oocytes collected from the experiment reported in A at indicated times, using antibodies directed against the proteins indicated on the right. Molecular weights (kDa) are indicated on the left. eNEBD and eMII: Cip1-injected oocytes collected at 6.5 hrs (NEBD50%) and 9.5 hrs (equivalent MII) respectively. C. *In vitro* Cdk1 kinase assay of oocytes from the experiment reported in A and B. Western blot analysis revealing the Cdk1 substrate (GST-PP1) and its phosphorylated form at T320 (GST-pT320-PP1).

The Cip1-injected oocytes do not show any morphological sign of NEBD and they do not progress to MI and MII. Therefore, to compare Cip1-injected oocytes and control oocytes at NEBD or in MII, Cip1-injected oocytes were collected at the time when 50% of the control oocytes reach NEBD (NEBD equivalent time, eNEBD) or 3 hrs later (MII equivalent time, eMII).

### Cyclin B1 accumulation is an early event, while Mos accumulation requires Cdk1 activity

During meiotic maturation, extensive changes in protein abundance take place. Two proteins have been reported to accumulate during the oocyte G2/M transition, Mos and Cyclin B1 (Figs. 1 and 3A) [13– 15,18,28]. Interestingly, the suppression of their translation delays Cdk1 activation, suggesting that their accumulation regulates Cdk1 activation [13,28]. Since Cdk1 activation occurs into two steps, their accumulation could be required for the formation of the starting Cdk1 activity or be involved in the autoamplification loop to convert pre-MPF into MPF. In the first model, their accumulation would occur independently of the initial activation of Cdk1 activity, while in the second model, the accumulation would take place only downstream the starting amount of Cdk1. To address this issue, we evaluated the accumulation of these two proteins after exposure to progesterone in oocytes where the activation of starting Cdk1 has been prevented by Cip1 injection. Cyclin B1 accumulation took place in response to progesterone in Cip1-injected oocytes (Fig. 3A-B). The level of Cyclin B1 at NEBD or eNEBD was similar in the presence (1.0 ± 0.10) or absence (0.98 ± 0.10) of Cdk1 activity (Fig. 3B). The level of the protein increased further in MII in control oocytes (1.9 ± 0.20) in contrast to Cip1-injected ones (1.3 ± 0.10) (Fig. 3A-B). Therefore, two distinct mechanisms control Cyclin B1 accumulation during the G2/M transition: Cyclin B1 begins to accumulate independently of Cdk1 activity; then, once Cdk1 is fully activated at NEBD, its accumulation continues in a Cdk1-dependent manner.

**Fig. 3.**
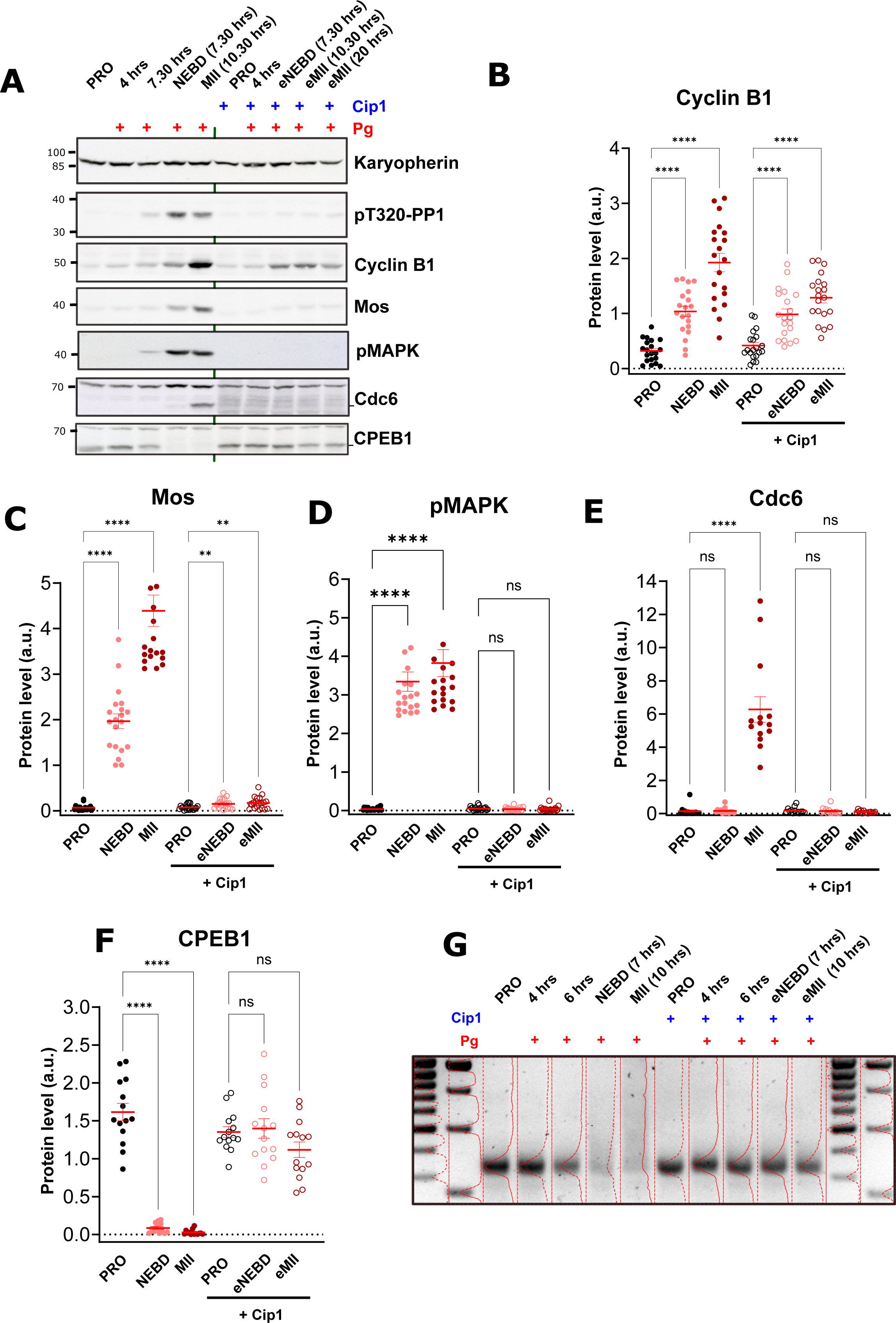
The level of key proteins is differentially regulated, either dependently or independently of Cdk1. A. Oocytes were injected or not with Cip1. After overnight incubation, progesterone (Pg) was added. Oocytes were collected at the indicated times and extracts were analyzed by western blot using antibodies directed against the proteins indicated on the right. Molecular weights (kDa) are indicated on the left. eNEBD and eMII: Cip1-injected oocytes collected at time of NEBD (eNEBD: equivalent NEBD) or MII (eMII: equivalent MII) in Pg control oocytes. **B-F.** Cyclin B1 (B), Mos (C), pMAPK (D), Cdc6 (E) and CPEB1 (F) signals were quantified from oocytes injected (empty dots) or not (filled dots) with Cip1 and collected at prophase (PRO), NEBD, metaphase II (MII) or equivalents times for Cip1-injected oocytes (PRO, eNEBD, eMII). 20 (for Cyclin B1, Mos and pMAPK) or 14 (for Cdc6 and CPEB1) biological replicates were quantified. One-way ANOVAs test with multiple comparisons were performed to evaluate the statistical significance. **: p<0.01, ****: p<0.0001, ns: not significant. G. Oocytes were injected or not with Cip1. After overnight incubation, progesterone (Pg) was added. Oocytes were collected at the indicated times, at NEBD, metaphase II (MII) or equivalents times for Cip1-injected oocytes (PRO, eNEBD, eMII). The analysis of the poly(A) tail length of the endogenous Cyclin B1 mRNA was performed by PAT-assay.

The Mos/MAPK activation was also proposed to control the activation of Cdk1 [18,28]. Mos is detectable before NEBD at the same time as the formation of the starting activity of Cdk1 (Fig.1A and Fig. 3A), making it difficult to determine if it is involved in activating the starting activity of Cdk1 or in the autoamplification loop. To address this question, Cip1 was injected in oocytes that were then stimulated by progesterone. Cip1 injection almost abolished the accumulation of Mos occurring in response of progesterone (Fig. 3A and 3C). A statistically significant 2-fold increase of Mos was observed in Cip1-injected oocytes in response to progesterone (from 0.08 ± 0.01 in PRO to 0.15 ± 0.02 in eNEBD), but this increase is very modest compared with the 25-fold increase occurring in control oocytes (from 0.07 ± 0.01 in PRO to 1.97 ± 0.16 at NEBD) (Fig. 3C). The 2-fold increase in the level of Mos in Cip1-injected oocytes was unable to activate MAPK, as demonstrated by the complete absence of the phosphorylated Erk1/2 (Fig 3A and D). Hence, the activation of the Mos/MAPK pathway does not occur in response to progesterone when Cdk1 activation is prevented by Cip1, showing that it depends on the formation of the starting activity of Cdk1. Therefore, the implication of Mos/MAPK in Cdk1 activation lies at the level of the auto-amplification loop, explaining the delay of NEBD observed when Mos translation is impaired [28].

Another regulator of Cdk1 activity during meiotic divisions is the DNA pre-replication component Cdc6 [26]. Cdc6 accumulates in an highly regulated manner during meiotic maturation and becomes detectable by western blot after NEBD (Fig. 1) [26,29,30]. We have previously shown that the suppression of the Mos/MAPK module accelerates Cdc6 accumulation, demonstrating that this pathway negatively regulates Cdc6 accumulation under physiological conditions [26]. We determined the pattern of Cdc6 accumulation in oocytes injected with Cip1, where both the activation of Cdk1 and Mos/MAPK are prevented. Cdc6 accumulation did not occur in Cip1-injected oocytes that were stimulated by progesterone (Fig. 3A and 3E). Therefore, Cdc6 accumulation is a “late event” that does not participate in the control of Cdk1 activation at the G2/M transition.

One of the master regulators of translation in oocytes is CPEB1 [31,32]. CPEB1 degradation was shown to allow Cyclin B1 polyadenylation during the G2/M transition in *Xenopus* oocytes, an event that is critical for mRNA translation [33]. Since Cyclin B1 accumulates independently of Cdk1 activation (Fig. 3A-B) and CPEB1 degradation is required for Cyclin B1 polyadenylation/translation, we expected CPEB1 to be degraded as an “early event”, independently of Cdk1 activation, to promote the early period of Cyclin B1 accumulation. However, as shown in Fig. 1, CPEB1 degradation begins in parallel to the first activation of Cdk1. To determine whether its degradation depends on Cdk1 activity, Cdk1 activation was prevented by Cip1 injection. Interestingly, CPEB1 was stable in Cip1-injected oocytes stimulated by progesterone (Fig. 3A and F). Therefore, CPEB1 degradation is a “late event” that cannot account for the early accumulation of Cyclin B1. Since CPEB1 degradation is known to be required for Cyclin B1 translation, we ascertained the polyadenylation status of endogenous Cyclin B1 mRNAs with a PAT-assay. Surprisingly, Cyclin B1 mRNA was not polyadenylated in oocytes where Cdk1 activation was prevented by Cip1 injection (Fig. 3G). This demonstrates that the early period of Cyclin B1 accumulation is not controlled by mRNA polyadenylation, which occurs only downstream Cdk1 activation.

### Cyclin B1 accumulation does not depend on the stimulation of its translation

We have shown that Cyclin B1 starts to accumulate independently of Cdk1 activation and CPEB1 degradation and continues to accumulate at NEBD in a Cdk1-dependent manner (Fig. 3A-B). Cyclin B1 accumulation can be due to an increase in the translation rate or to a decrease in the degradation of the protein. To address this issue, we designed a translation reporter to monitor Cyclin B1 translation independently of the turnover of the protein. The 5’- and 3’-UTRs of Cyclin B1 were cloned upstream and downstream of an open reading frame encoding a GST-v5 recombinant protein (Fig. 4A). In such a way, the translation of the GST-v5 reporter is controlled as the translation of the endogenous Cyclin B1. This reporter was co-injected with a constitutive probe encoding for a MBP-GST-v5 fusion protein without any UTRs except for a poly(A) tail, guaranteeing constant accumulation of the MBP-GST-v5 protein and acting as a control (Fig. 4A). After overnight incubation, oocytes were incubated with progesterone to trigger meiosis resumption. During the overnight incubation, the control polyadenylated probe was highly translated and the MBP-GST-v5 accumulated, while the Cyclin B1 reporter was only moderately translated (Fig. 4B). Stimulation by progesterone did not modify the translation rate of the control constitutive probe. In contrast, the translation rate of the Cyclin B1 reporter increased (Fig. 4B-C). It should be noted that the GST-v5 protein undergoes an electrophoretic retardation, and that consequently the unshifted and shifted bands were used for the reporter quantifications. Interestingly, when Cdk1 activation was prevented by Cip1-injection, no increase in the rate of translation of the Cyclin B1 reporter was detected (Fig. 4B-C). This result shows that the early accumulation of Cyclin B1 is not due to an increase in the mRNA translation. This result is consistent with our observation that the early accumulation of Cyclin B1 occurs in the absence of CPEB1 degradation and mRNA polyadenylation (Fig. 3F-G).

**Fig. 4.**
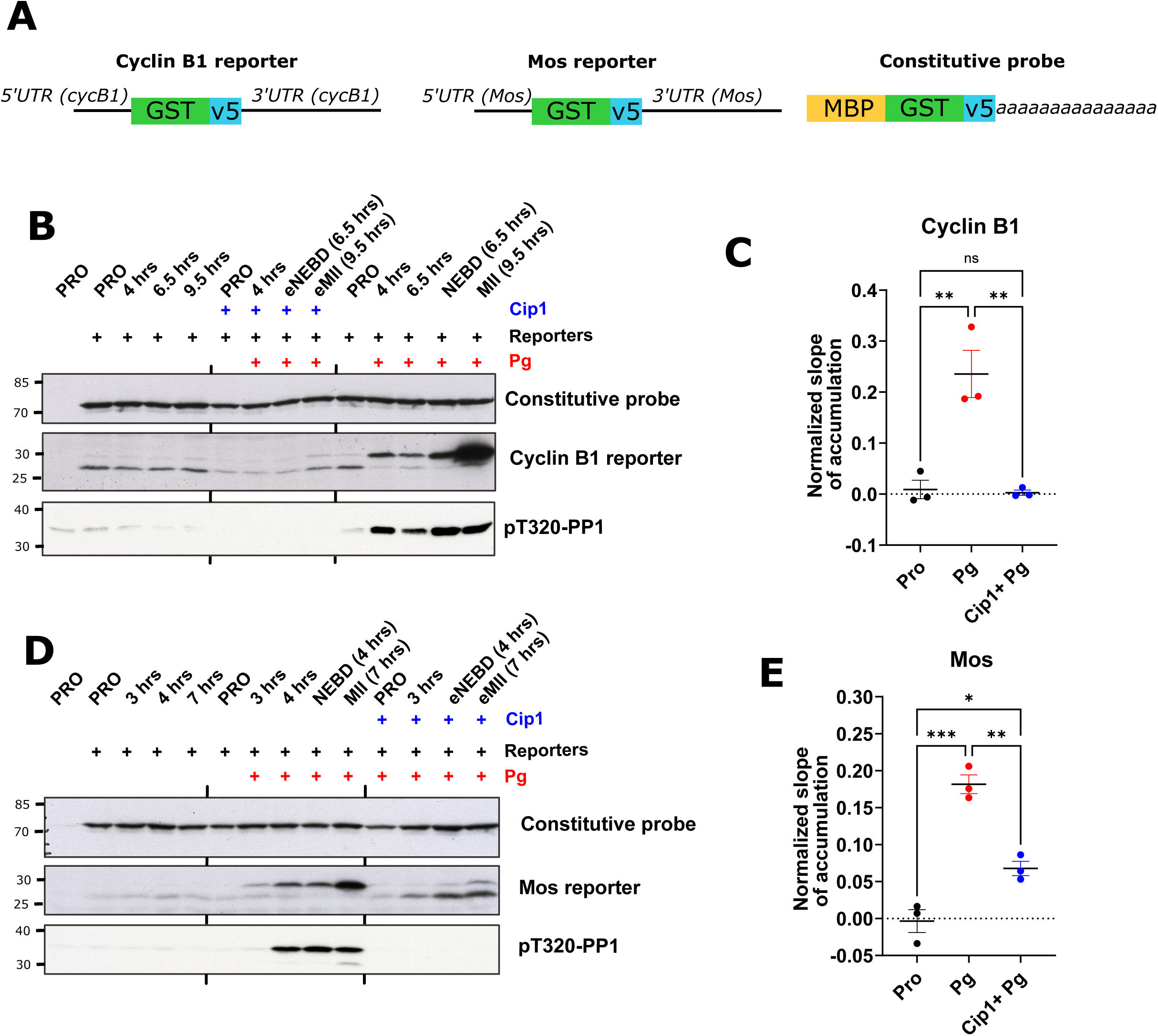
Mos translation increases independently of Cdk1 activity, while the activation of Cyclin B1 translation requires the activation of Cdk1. A. Schematic representation of the translation reporters used to monitor Cyclin B1 and Mos translation. B. Oocytes were injected or not with Cip1. One hour later, they were injected with a 5:1 mix of the Cyclin B1 reporter and the constitutive probe displayed in A. After overnight incubation, progesterone (Pg) was added or not. Oocytes were collected at prophase (PRO), at the indicated times after Pg addition, at NEBD, metaphase II (MII) or the equivalent times (eNEBD, eMII) for the Cip1-injected oocytes. Extracts were analyzed by western blot with antibodies against V5 (two top panels) and pT320-PP1 (lower panel). Molecular weights (kDa) are indicated on the left. C. The slopes of translation for prophase oocytes (PRO), oocytes incubated with progesterone (Pg) and Cip1-injected oocytes incubated with progesterone (Cip1+Pg) from three biological replicates of the experiment described in B were plotted. One-way ANOVAs test with multiple comparison were performed to evaluate the statistical significance. **: p<0.01, ns: not significant. D. Oocytes were injected or not with Cip1. One hour later, they were injected with a 5:1 mix of the Mos reporter and the constitutive probe displayed in A. After overnight incubation, progesterone (Pg) was added or not. Oocytes were collected at prophase (PRO), at the indicated times after Pg addition, at NEBD, metaphase II (MII) or the equivalent times (eNEBD, eMII) for the Cip1-injected oocytes. Extracts were analyzed by western blot with antibodies against V5 (two top panels) and pT320-PP1 (lower panel). Molecular weights (kDa) are indicated on the left. E. The slopes of translation for prophase oocytes (PRO), oocytes incubated with progesterone (Pg) and Cip1-injected oocytes incubated with progesterone (Cip1+Pg) from three biological replicates of the experiment described in D were plotted. One-way ANOVAs test with multiple comparison were performed to evaluate the statistical significance. *: p<0.05, ***: p<0.001.

### The increase in Mos translation is not sufficient to accumulate the protein

It was shown that Mos mRNA polyadenylation is activated in response of progesterone independently of CPEB1 degradation, suggesting it could occurs independently of Cdk1 activation [33]. To get further insight in the regulation of Mos translation, we evaluated the translation pattern of Mos during meiotic maturation by using a translation reporter similar to the Cyclin B1 one (Fig. 4A). The Mos reporter translation was extremely low in prophase oocytes during the overnight incubation, as compared to the polyadenylated control construct (Fig. 4D). The Mos reporter translation increased in response to progesterone before NEBD (Fig. 4D). Interestingly, the activation of Mos reporter translation also took place when Cdk1 activation was prevented by Cip1 injection, even though not reaching the same level than in the control oocytes (0.18 ± 0.01 in control oocytes versus 0.07 ± 0.01 in Cip1-injected oocytes) (Fig. 4D-E). Therefore, the translation of Mos begins before and independently of Cdk1 activation and is further enhanced by the activation of Cdk1. However, the protein does not accumulate (Fig. 3A and C), suggesting that newly translated Mos protein is not stable due to an active degradation. Interestingly, it was shown that Mos is stabilized by the phosphorylation at S3 by Cdk1 [34]. This could explain that Mos accumulation is only detected downstream the activation of the starter Cdk1 (Fig. 3A and C) despite an active translation occurring earlier (Fig. 4D-E).

### Two waves of protein translation occur during the *Xenopus* G2/M transition

The changes in protein abundance during *Xenopus* oocyte meiotic divisions are regulated at the post-transcriptional level since transcription is completely silent from the end of oocyte growth until the mid-blastula transition of the embryo (13^th^ cell division cycle in *Xenopus*)[35]. Indeed, the translation of new proteins is a critical step regulating meiotic divisions in vertebrates and many invertebrates [11,36–38]. In all vertebrates, excluding the small rodents, the *de novo* translation of proteins is required for the oocyte G2/M transition and its inhibition by cycloheximide prevents NEBD in response to either progesterone or PKI [9,37]. According to these results, protein translation should occur as an “early event”, as we have shown for Mos (Fig. 4D-E). However, while cycloheximide completely blocks Cdk1 activation, the suppression of Mos translation only delays the G2/M transition[28], suggesting that other mRNAs need to be translated to activate Cdk1. We measured the global level of protein translation in *Xenopus* oocytes using the SUnSET method to determine the magnitude of the translation in prophase oocytes and during meiotic divisions, before and after Cdk1 activation [39]. This technique allows to measure the incorporation of puromycin into newly synthesized peptides, giving a readout of the overall activity of the ribosomes. A progressive increase of puromycin incorporation in proteins was detected in oocytes after progesterone stimulation reaching its maximum at NEBD (from 0.80 ±0.14 in PRO to 1.78±0.19 in NEBD) (Fig. 5). Interestingly, the rate of translation increased in response to progesterone even if the activation of Cdk1 was prevented by Cip1 injection (from 0.52 ±0.08 in PRO to 1.30±0.12 at eNEBD), although it did not reach the level of the control oocytes (Fig. 5). These results show that the activation of translation is both an “early” and a “late event”, the early period occurring independently of Cdk1 activation. Hence, distinct pathways control translation during meiotic divisions.

**Fig. 5.**
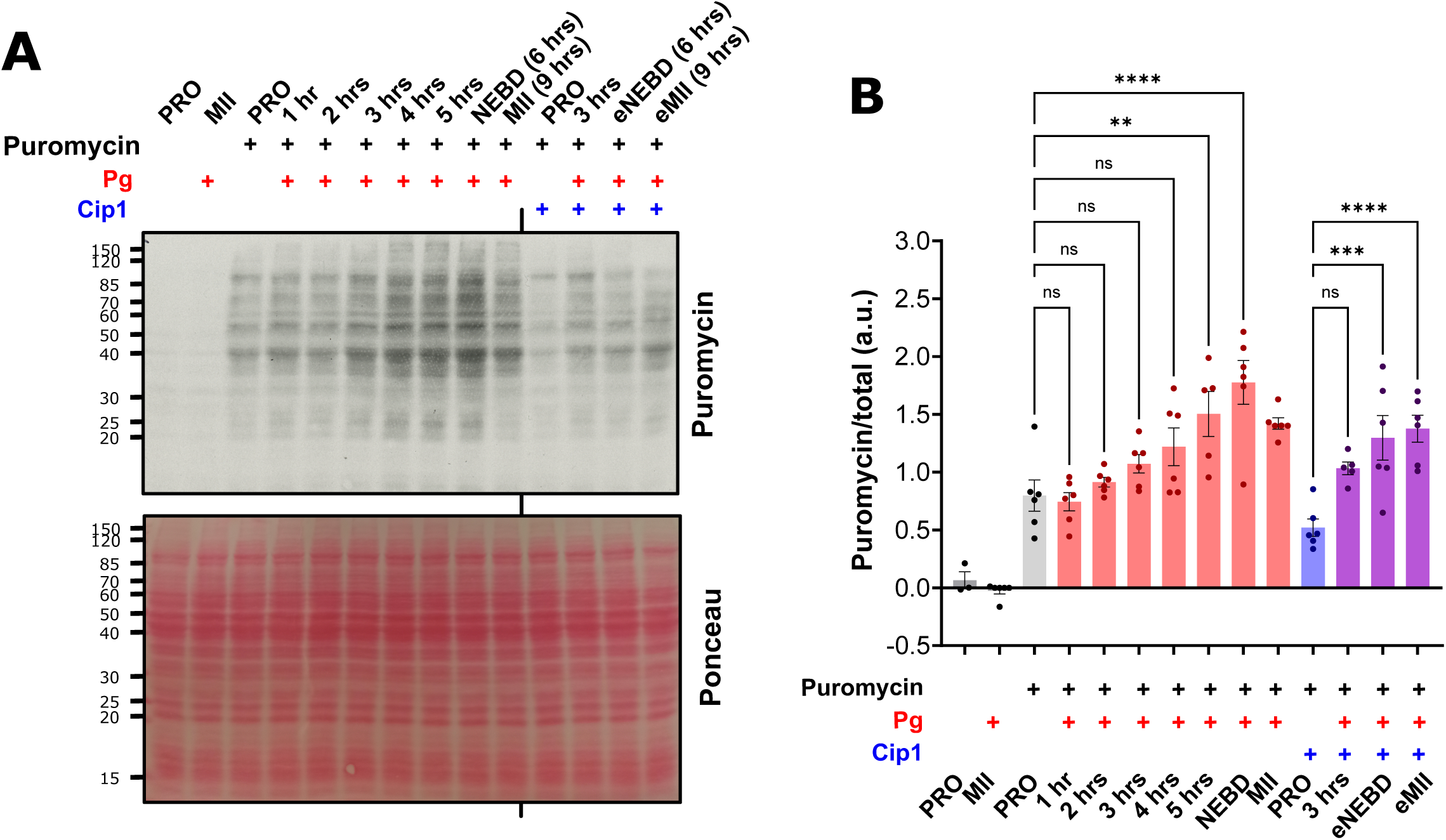
Two waves of protein translation take place during meiosis resumption, the early one occurring independently of Cdk1-activation. A. Oocytes were injected or not with Cip1. After overnight incubation, progesterone (Pg) was added. 10 min before collection, oocytes were injected with puromycin. Oocytes were collected at the indicated times. eNEBD and eMII: Cip1-injected oocytes collected at time of NEBD (eNEBD: equivalent NEBD) or MII (eMII: equivalent MII) in Pg control oocytes. The incorporation of puromycin into the *de novo* synthesized peptides was visualized by western blot. The total amount of proteins was visualized by Ponceau Red staining. Molecular weights (kDa) are indicated on the left. B. Quantification of 6 different biological replicates of the experiment described in A. The puromycin signals were normalized on the signals of total amount of proteins visualized by Ponceau Red staining. **: p<0.01, ***: p<0.001, ****: p<0.0001, ns: not significant.

### PKA inhibition is sufficient to promote meiotic maturation through the activation of the early events

Our aim is to understand which are the molecular events triggered by PKA downregulation and controlling Cdk1 activation. Therefore, it was important to ascertain that the early events identified in Cip1-injected oocytes in response to progesterone are reproduced by a specific inhibition of PKA, in the absence of progesterone. For this purpose, oocytes were injected with PKI, and then injected with Cip1 or not. As already reported [6,8], PKI injection induced meiotic maturation and promoted Cdk1 and MAPK activation (Fig. 6A). Both events were fully blocked by Cip1 injection (Fig. 6A). All the associated biochemical events as the accumulation of Cyclin B1, Mos and Cdc6, CPEB1 degradation, activation of protein translation and Cdk1 activation, were induced by PKI injection similarly to progesterone (Fig. 6B-C). Importantly, Cyclin B1 accumulation induced by PKI was not blocked by Cip1-injection whereas Mos and Cdc6 accumulation as well as CPEB1 degradation did not take place in the absence of Cdk1 activation (Fig. 6B). Hence, the biochemical events analysed in this paper are controlled identically by progesterone and its downstream target, PKA.

**Fig. 6.**
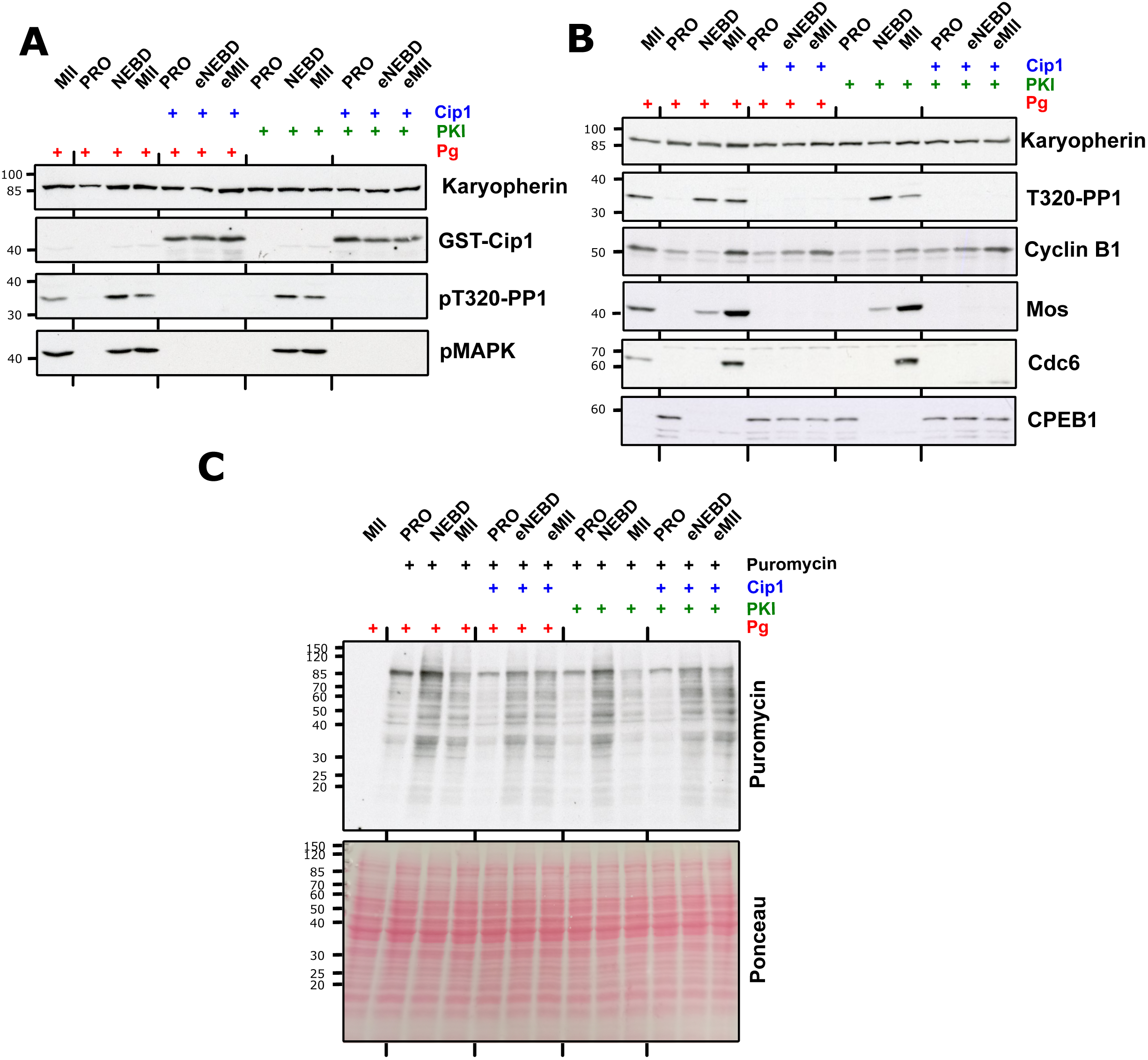
**PKI induces meiosis resumption by activating the early events.** Oocytes were injected or not with Cip1. After overnight incubation, progesterone (Pg) was added in the external medium or PKI was injected in the oocytes. Oocytes were collected at prophase (PRO), NEBD, metaphase II (MII) or the equivalent times (eNEBD, eMII) for the Cip1-injected oocytes. The experiment was performed twice with similar results. **A-B**. Extracts were analyzed by western blot using antibodies directed against the proteins indicated on the right. Molecular weights (kDa) are indicated on the left. C. 10 min before the time of collection, oocytes were injected with puromycin. The incorporation of puromycin into *de novo* synthesized peptides was visualized by western blot. The total amount of proteins was visualized by Ponceau Red staining. Molecular weight (kDa) are indicated on the left.

### Arpp19 control neither Cyclin B1 accumulation nor the Cdk1-Cyclin B complexes

We have identified three events that depend on the inactivation of PKA and occur before and independently of Cdk1 activation: Cyclin B1 accumulation, Mos translation and the increase in the global rate of translation. Since these events depend on PKA inactivation, the dephosphorylation of PKA substrates should promote directly or indirectly these “early events”. Up to date, a single substrate of PKA controlling *Xenopus* meiosis resumption, Arpp19, has been identified [19]. Arpp19 is phosphorylated on S109 by PKA and dephosphorylated by PP2A-B55 in response to progesterone [20]. A phosphomimetic mutant, Arpp19-S109D, blocks Cdk1 activation induced by progesterone [19]. However, the molecular mechanism by which PKA-phosphorylated Arpp19 inhibits meiosis resumption is still unknown. Therefore, we tested whether the phosphomimic Arpp19 mutant could impair any of the “early events” we identified. Oocytes were injected with Arpp19-S109D and, after an overnight incubation, were stimulated by progesterone. As expected, the S109D-phosphomutant suppressed the oocyte G2/M transition as shown by the absence of the increase in PP1 phosphorylation at T320 and the absence of Cdk1 activity assessed in an *in vitro* kinase assay (Fig. 7A-B).

**Fig. 7.**
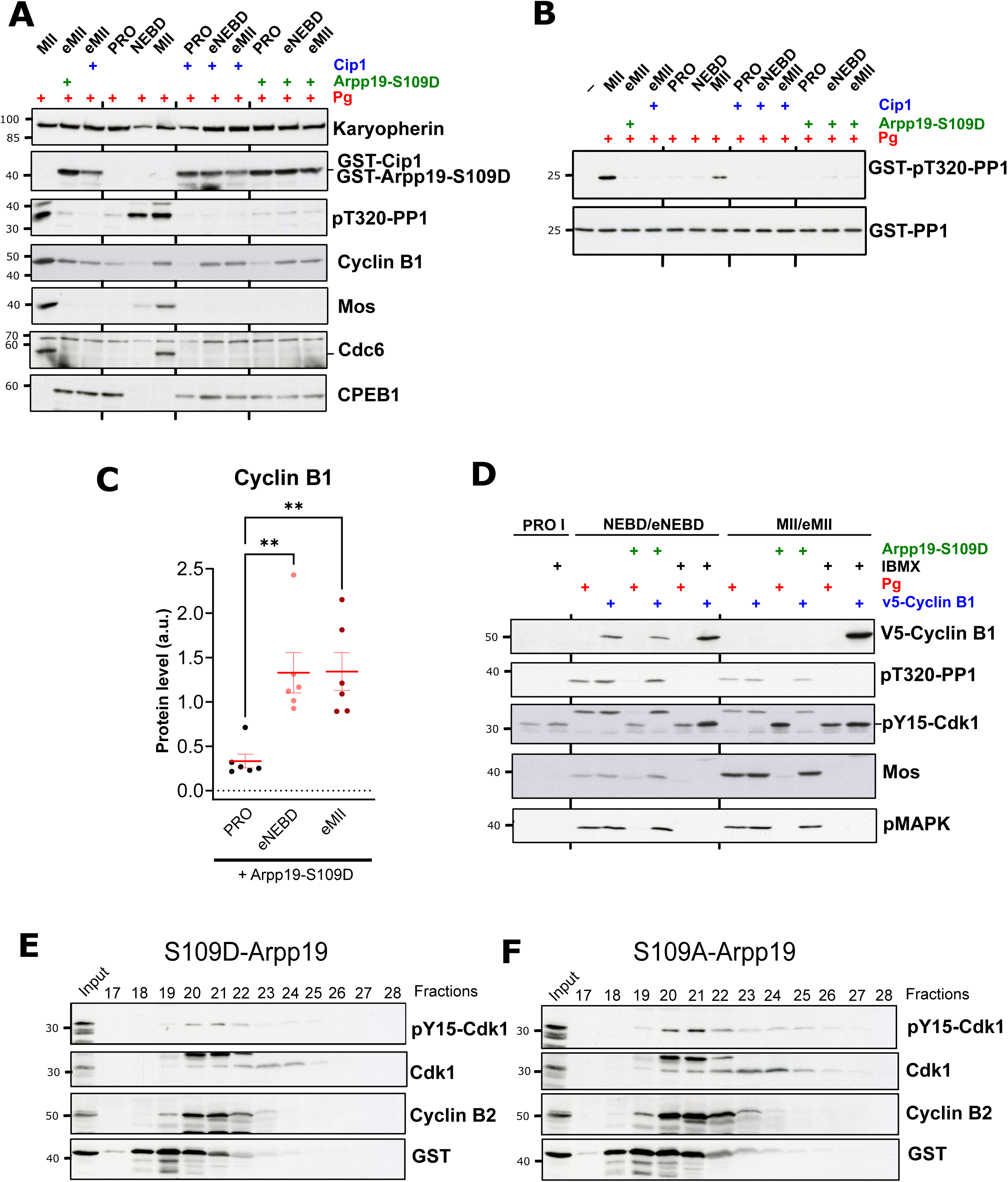
Arpp19-S109D does not prevent the accumulation of Cyclin B1 or Cdk1-Cyclin B activity. A. Oocytes were injected or not with either Cip1 or Arpp19-S109D. After overnight incubation, progesterone (Pg) was added in the external medium. Oocytes were collected at prophase (PRO), NEBD, metaphase II (MII) or the equivalent times (eNEBD, eMII) for the Cip1- and Arpp19-S109D injected oocytes. Extracts were analyzed by western blot using antibodies directed against the proteins indicated on the right. Molecular weights (kDa) are indicated on the left. Note that recombinant GST-Cip1 and GST-Arpp19- S109D proteins have the same molecular weight. B. *In vitro* Cdk1 kinase assay of oocytes from the experiment reported in A. Western blot analysis revealing the Cdk1 substrate (GST-PP1) and its phosphorylated form at T320 (GST-pT320-PP1). Molecular weight (kDa) is indicated on the left. C. Quantification of the levels of Cyclin B1 of GST-Arpp19-S109D-injected oocytes obtained as described in A. 6 independent biological replicates were analyzed. One-way ANOVAs test with multiple comparison were performed to evaluate the statistical significance. **: p<0.01. D. Oocytes were injected or not with Arpp19-S109D. After overnight incubation, the oocytes were incubated or not in the presence of 1mM IBMX. One hour later, oocytes were incubated with progesterone (Pg) or injected with mRNA encoding v5-Cyclin B1. Oocytes were collected at prophase (PRO), NEBD, metaphase II (MII) or the equivalent times (eNEBD, eMII) for the Arpp19-S109D- or IBMX-treated oocytes. Extracts were analyzed by western blot using antibodies directed against the proteins indicated on the right. Molecular weights (kDa) are indicated on the left. **E-F**. Oocytes were injected with Arpp19-S109D (**E**) or Arpp19-S109A (**F**). The protein extracts corresponding to 50 injected oocytes were fractionated on a Superose 6 column. The fractions were analyzed by western blot using antibodies directed against the proteins indicated on the right. Molecular weights (kDa) are indicated on the left.

We determined whether the early accumulation of Cyclin B1 depends on the phosphorylation status of Arpp19. Oocytes were injected with Arpp19-S109D and, after an overnight incubation, were stimulated by progesterone. Importantly, the early accumulation of Cyclin B1 occurred in response to progesterone in the presence of Arpp19-S109D, as in Cip1-injected oocytes (Fig. 7A and C). The “late events” that depend on Cdk1 activation were inhibited, as expected (Supplementary Figure). Therefore, Arpp19 is not the PKA substrate involved in the control of Cyclin B1 accumulation.

Although PKA-phosphorylated Arpp19 does not prevent Cyclin B1 accumulation induced by progesterone, it blocks Cdk1 activation. The phosphorylated form of Arpp19 could block Cdk1 activation by either inactivating the newly formed Cdk1-Cyclin B1 complexes that work as a starter Cdk1 activity, or by destabilizing the stockpiled pre-MPF, that is made of Cdk1 and Cyclin B2.

To test the first hypothesis, we determined whether Arpp19-S109D can block Cdk1 activation when induced by Cyclin B1 translation. Since Cyclin B1 translation occurs only downstream Cdk1 activation in oocytes (Fig. 4B-C), we bypassed the repressor signals present in its UTRs by using a new construct: a mRNA encoding Cyclin B1 where the UTRs have been substituted by a long poly(A) tail. The injection of this polyadenylated Cyclin B1 mRNA induced meiosis resumption in the absence of the hormonal stimulation, as seen by the activation of MAPK, the phosphorylation of PP1 at T320, and the dephosphorylation of Cdk1-Cyclin B complexes on Y15 (Fig. 7D). This shows that Cyclin B1 that is efficiently translated upon its injection in prophase oocytes, forms active complexes with Cdk1. Note that at the MII stage, the ectopic Cyclin B1 is no longer detected (Fig. 7D), because the translation of the *in vitro* polyadenylated-mRNAs decays over time and that the protein is degraded by the APC during the MI-MII transition. Importantly, the newly translated Cyclin B1 induced the activation of Cdk1 even in the presence of Arpp19-S109D (Fig. 7D). This result demonstrates that PKA-phosphorylated Arpp19 does not inhibit the newly formed Cdk1-Cyclin B1 complexes (Fig. 7D). Additionally, we compared the effects of constitutively active PKA with those of its phosphorylated substrate, Arpp19-S109D. Incubation of oocytes with the phosphodiesterase inhibitor IBMX prevents meiotic maturation induced by progesterone by maintaining high cAMP levels and hence high PKA activity [7,40] (Fig. 7D). Interestingly, IBMX treatment completely suppressed the ability of Cyclin B1 mRNA to activate Cdk1, a different behaviour than Arpp19-S109D (Fig. 7). This result suggests that additional PKA effectors exist in addition to Arpp19 that, under their phosphorylated form, prevent meiosis resumption by blocking the activation of newly Cdk1-Cyclin B1 complexes.

We then tested the second hypothesis by assessing if Arpp19-S109D could induce the dissociation of the already formed Cdk1-Cyclin B2 complexes (pre-MPF) in prophase oocytes. Oocytes were injected with Arpp19-S109D, then lysed and the oocyte extracts were fractionated by size-exclusion chromatography. We analysed the elution profile of Cyclin B2 and Cdk1 in the fractions to evaluate the integrity of the pre-MPF. Cyclin B2 and Y15-phosphorylated Cdk1 were eluted in fractions 20-21-22, while free-Cdk1 (not associated with Cyclin B2) was eluted in fractions 23 and 24 (Fig. 7E). Another Arpp19 mutant, Arpp19- S109A, whose injection does not affect Cdk1 activation and meiosis resumption [19], was used as a control. The same elution profile was obtained by fractionating oocytes injected with Arpp19-S109A (Fig. 7F).

Therefore, Arpp19-S109D does not supress the activation of Cdk1 neither by interfering with the accumulation of Cyclin B1, nor by inhibiting the newly formed Cdk1-Cyclin B1 complexes, nor by destabilizing the pre-MPF stockpiled in the oocyte.

### Arpp19 does not control the global rate of protein translation or the translation of Mos

Interestingly, the yeast homologs of Arpp19, igo1 and igo2, control the activation of the translational program at the exit from the quiescent state [41,42]. Since protein translation is activated before and independently of Cdk1 (Fig. 5) and is necessary for the G2/M transition, we evaluated whether Arpp19- S109D represses the early translation using the SUnSET approach. Oocytes were injected with Arpp19- S109D or Cip1 and after an overnight incubation, meiotic maturation was induced by progesterone. The injection of either Cip1 or Arpp19-S109D did not block the early wave of translation as seen by an increase of the level of protein translation at eNEBD and eMII (in Arpp19-S109D injected oocytes: from 0.77 ± 0.09 in PRO to 1.81 ± 0.22 at eNEBD and 1.70 ± 0.27 in eMII) (Fig. 8A-B).

**Fig. 8.**
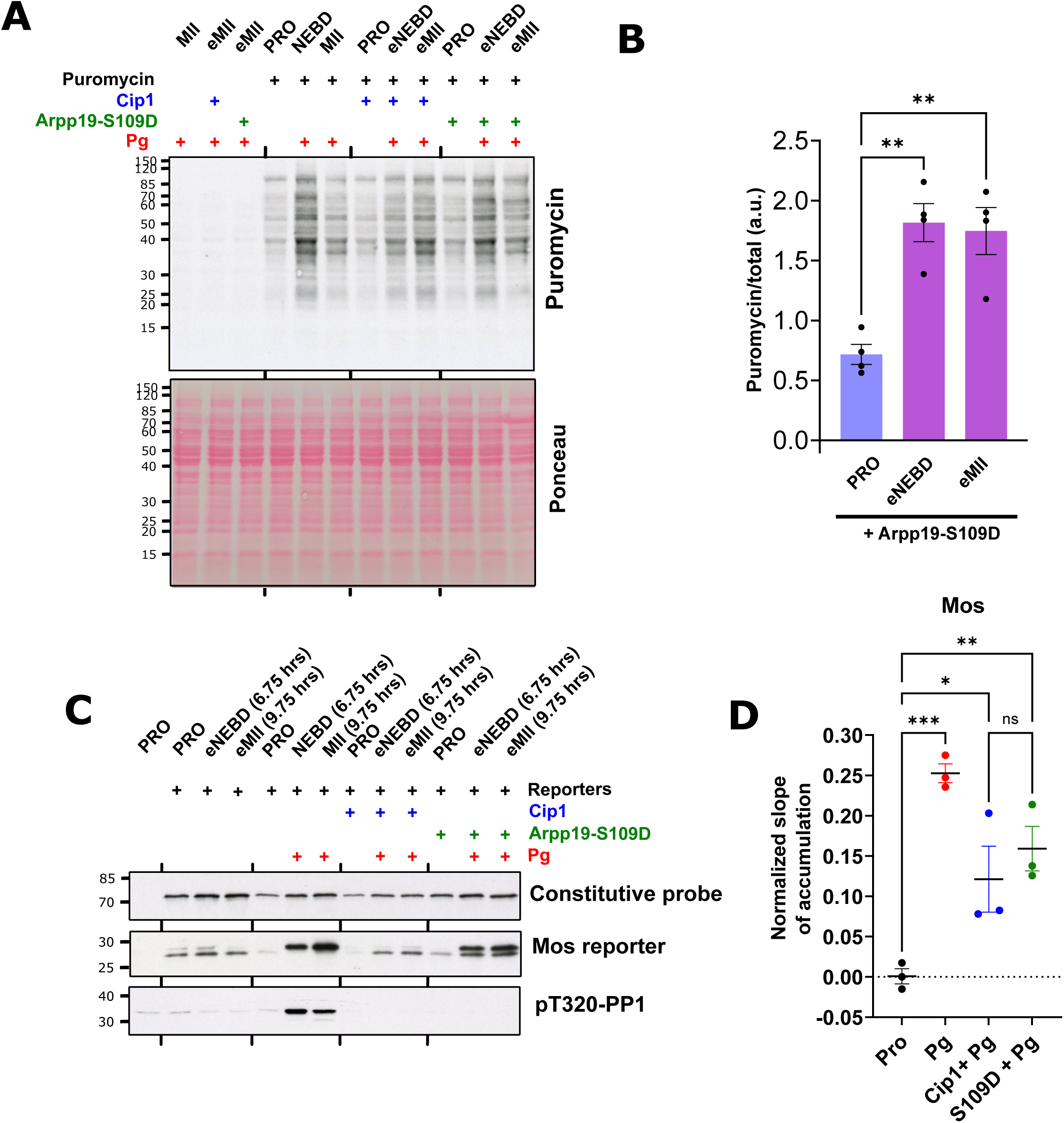
Arpp19-S109D does not block the global activation of protein translation or the translation of Mos. A. Oocytes were injected or not with either Cip1 or Arpp19-S109D. After overnight incubation, progesterone (Pg) was added. 10 min before the collection, oocytes were injected with puromycin. Oocytes were collected at prophase (PRO), NEBD, metaphase II (MII) or the equivalent times (eNEBD, eMII) for the Cip1- and Arpp19-S109D injected oocytes. The incorporation of puromycin into *de novo* synthesized peptides was visualized by western blot. The total amount of proteins was visualized by Ponceau Red staining. Molecular weights (kDa) are indicated on the left. B. Quantification of 4 biological replicates of the experiment described in A. The puromycin signals were normalized on the signals of total amount of proteins visualized by Ponceau Red staining. One-way ANOVAs test with multiple comparison were performed to evaluate the statistical significance. **: p<0.01. C. Oocytes were injected or not with either Cip1 or Arpp19-S109D. One hour later, they were injected with a 5:1 mix of the Mos reporter and the constitutive probe displayed in Fig. 4A. After overnight incubation, progesterone (Pg) was added or not in the external medium. Oocytes were collected at prophase (PRO), NEBD, metaphase II (MII) or the equivalent times (eNEBD, eMII) for the Cip1- and Arpp19-S109D injected oocytes. Extracts were analyzed by western blot with antibodies against V5 (two top panels) and pT320- PP1 (lower panel). Molecular weights (kDa) are indicated on the left. D. The slopes of translation for prophase oocytes (PRO), oocytes incubated with progesterone (Pg), Cip1-injected oocytes incubated with progesterone (Cip1+Pg) and Arpp19-S109D injected oocytes (S109D+Pg) from three biological replicates of the experiment described in C were plotted. One-way ANOVAs test with multiple comparison were performed to evaluate the statistical significance. *: p<0.05, **: p<0.01, ***: p<0.001, ns: not significant.

We have shown that Mos is among the early mRNAs whose translation increases independently of Cdk1 activation (Fig. 4D-E), even though Mos protein cannot accumulate before Cdk1 activation (Fig. 3A and C). We tested whether Arpp19-S109D affects the early translation of Mos using the translation reporter approach (Fig. 4A). Arpp19-S109D did not block the activation of Mos translation estimated by the Mos translation reporter (Fig. 8C-D). The levels of translation in response to progesterone, in the presence of either Arpp19-S109D or Cip1, were similarly increased, although less than in control oocytes (Fig. 8C-D).

Therefore, Arpp19-S109D does not inhibit Cdk1 activation by affecting the global activation of mRNA translation or by regulating Mos translation.

## DISCUSSION

Maintenance and release of the prophase arrest of oocytes play a critical role in sexual reproduction since they lead to the production of a fertilizable gamete. Two opposing kinases orchestrate these processes: first, PKA activity that maintains the prophase arrest in all vertebrate oocytes and whose inhibition is necessary for meiotic resumption; second, Cdk1 activity whose activation is promoted by PKA inhibition and necessary for oocyte division. Correct regulation of the multi-steps pathways connecting PKA inhibition and Cdk1 activation is of prime importance for the success of the female meiotic divisions. However, these pathways are far from being known. They involve several PKA substrates, of which only one is currently identified [19], and protein translation, whose inhibition blocks Cdk1 activation [9]. Moreover, specific proteins accumulate, as Cyclin B1 and Mos, and play a role in Cdk1 activation [3,13,28]. However, it is difficult to determine which events controlled by PKA inhibition are necessary for Cdk1 activation. Indeed, basic Cdk1 activation takes place before NEBD. The appearance of the white spot signaling NEBD is the only marker that can be used to study meiotic divisions in the amphibian oocyte. But when this spot appears, Cdk1 is already partly active.

In this work, we used a direct and specific inhibitor of Cdk1, Cip1, to isolate the events initiated by PKA inhibition independently of Cdk1, and potentially responsible for its activation. We thus discovered three events directly dependent on PKA inhibition, which we call “early events”, none of which being under the control of the only identified PKA substrate regulating Cdk1 activation, Arpp19.

Our results show that the regulation of protein accumulation induced by PKA inactivation is more complex than previously thought and involves the control of both the rate of synthesis and degradation of proteins. We have shown that Mos translation is initiated in response of progesterone independently of Cdk1 activity, but the protein does not accumulate until Cdk1 activation. The E3-ubiquitine ligase controlling Mos degradation remains unknown, however the molecular connection between Mos stability and Cdk1 could rely on the ability of Cdk1 to phosphorylate and to stabilize Mos during meiosis II [34]. Surprisingly, Cyclin B1 is regulated in the opposite manner. It accumulates in response to progesterone independently of Cdk1 activation [14], but the rate of Cyclin B1 translation is not increased in the absence of Cdk1 activity. Moreover, the Cyclin B1 accumulation taking place in Cip1-injected oocytes does not require CPEB1 degradation and polyadenylation of its mRNA. These results highlight that Cyclin B1 stability, but not its translation, is regulated by PKA downregulation. The mechanism regulating Cyclin B1 turnover during the prophase arrest and early meiosis resumption is not known in *Xenopus* oocytes. In mouse oocytes, it was shown that Cyclin B1 turnover is regulated by the APC/Cdh1 ubiquitin ligase. Indeed, Cdh1 knockdown induces meiosis resumption in the presence of high PKA activity [43]. Surprisingly, Cdh1 inactivation by antisense injection inhibits meiosis resumption in *Xenopus* oocytes while overexpression of the protein facilitates the process [44]. Cdh1 may act in an APC-independent manner in *Xenopus* oocytes [44], explaining this apparent paradox. This pleiotropic function of Cdh1 makes it difficult to evaluate its impact on Cyclin B1 accumulation in *Xenopus* oocytes. The other regulator of APC, Cdc20, is responsible of Cyclin B degradation in Anaphase. However, Cdc20 depletion by antisense oligonucleotides does not affect the kinetics of meiotic maturation or the accumulation of Cyclin B [45]. The identification of the E3 ligases controlling the turnover of Mos and Cyclin B upstream Cdk1 activation should be a direction of future research.

Interestingly, both Cyclin B1 and Mos accumulation are regulated by two-steps mechanism (Fig. 9). The first step occurs before Cdk1 activation, when Cyclin B1 is stabilized and Mos translation is activated. The second one takes place downstream the starting amount of Cdk1 activity and involves the activation of Cyclin B1 translation and Mos stabilization. This intertwined regulation results in a positive feedback loop between Cdk1 activity and its regulators contributing to the timely and irreversible decision to go through the G2/M transition. Since we have shown that Mos accumulation and MAPK activation occur downstream the activation of the starter Cdk1, this limits the involvement of the Mos/MAPK module in Cdk1 activation to the second step, the auto-amplification loop. Upon the inhibition of the Mos/MAPK pathway, the starting amount of Cdk1 normally forms because of the progesterone-induced accumulation of Cyclin B1, but it takes longer to activate the auto-amplification loop required for NEBD. This model would explain the delay in NEBD observed in oocytes treated with morpholino against Mos or a pharmacological inhibitor of MAPK [13,28,46]. Conversely, when Cyclin B1 accumulation is blocked, the starting amount of Cdk1 cannot be formed by the recruitment of Cyclin B1. Hence, the oocytes rely only on the Mos/MAPK to activate Cdk1-Cyclin B2 complexes. Under these conditions, Mos translation that starts independently from Cdk1 activation would result in a slow accumulation of Mos due to the high turn-over of the protein. After several hours, a little pool of active MAPK could initiate the activation pre-MPF into a starting amount of Cdk1 activity. This activity would stabilize Mos, leading to the full activation of MAPK and an increased conversion of pre-MPF into MPF through the auto-amplification mechanism.

**Fig. 9:**
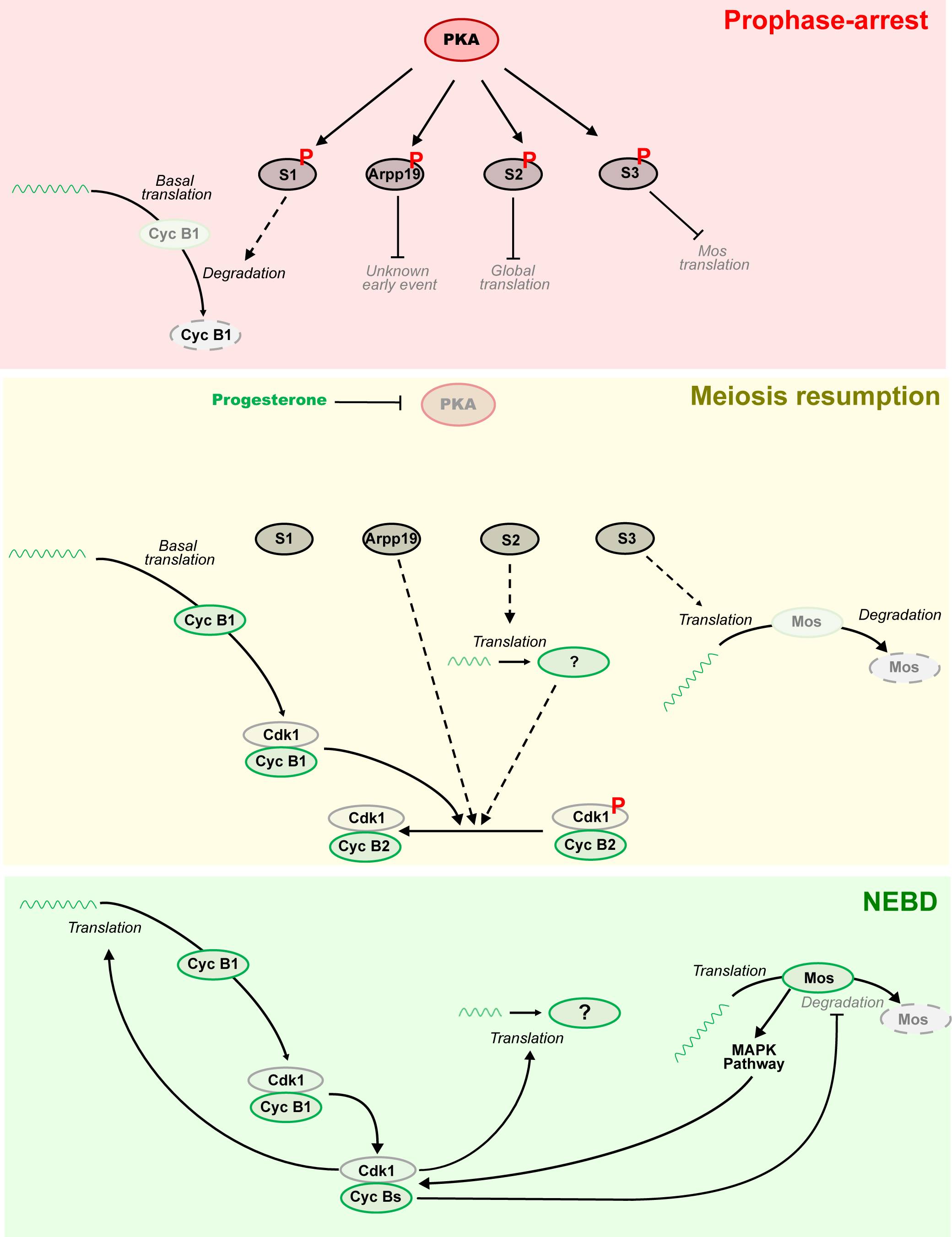
Multiple pathways control the activation of Cdk1 in *Xenopus* oocytes. **Prophase-arrest, upper panel**. The high activity of PKA and the phosphorylation of its substrates maintains the prophase arrest. PKA-phosphorylated substrates, S1, S2, S3 and Arpp19, control directly or indirectly the early eve, as the turnover of Cyclin B1 and the inhibition of the translation of early mRNAs, which includes the one encoding for Mos. **Meiosis resumption, middle panel**. In response to progesterone, PKA activity is inhibited resulting in dephosphorylation of its substrates. The accumulation of Cyclin B1 starts without an increase in the rate of its mRNA translation. The early wave of protein translation is also activated. This wave includes the mRNA encoding for Mos. Even though Mos translation increases, but the protein cannot accumulate. A starting amount of active Cdk1 is formed by the cooperation of two interconnected mechanisms: Cyclin B1 accumulation binds to free Cdk1; and *de novo* translation of proteins, including Mos, promotes the activation of a small portion of the pre-MPF formed of Cdk1-Cyclin B2. **NEBD, lower panel**. The starting amount of active Cdk1 promotes the auto-amplification loop, Cdk1 is fully activated due to the activation of stockpiled Cdk1-Cyclin B2 complexes. Full-Cdk1 activation promotes the translation of Cyclin B1, the late wave of translation, and the stabilization of Mos that accumulates activating the MAPK pathway.

It was shown more than 40 years ago that a strong increase in protein translation takes place at NEBD time in response to progesterone or injection of active MPF [47–49]. Indeed, the activation of protein translation during oocyte meiotic maturation involves hundreds of mRNAs, in mouse, *Xenopus* and human [37,38,50,51]. However, all these measurements were unable to discriminate the amplitude of translation activation taking place before or after Cdk1 activation. Our work shows that around half of the activation of protein translation occurring during the G2/M transition takes place independently of Cdk1 activation. Clearly, the early wave of translation does not only include Mos [15,28]. The identification of additional mRNAs composing this early wave of translation will be important in shedding light on how protein translation controls Cdk1 activation in oocytes, but potentially also meiotic divisions or early embryogenesis.

In addition to the early wave of translation, the translation of Mos and the accumulation of Cyclin B1 (Fig. 9), additional “early events” may exist. Recently, a partial degradation of the RNA-Binding protein, Zar1L, was observed within 2 hrs after progesterone exposure and was proposed to regulate Mos translation [52]. Zar1L degradation is a potential “early event” that could be tested using our Cip1-based experimental approach and could provide a molecular link to the early translation of Mos.

The discovery that multiple “early events” connect PKA downregulation to Cdk1 activation also suggests the existence of several PKA effectors that regulate them (Fig. 9). This model is supported by two lines of evidence. First, PKA inhibition by progesterone and PKI triggers the early events even in the presence of the phosphomimic mutant of the only known PKA effector, Arpp19. This result suggests that Arpp19 is not involved in the regulation of these events. Since these events depend on PKA inactivation, other PKA effectors must control them. Second, maintaining high PKA activity with IBMX blocks the ability of Cyclin B1 polyadenylated mRNAs to induce Cdk1 activation [53], whereas Arpp19-S109D is unable to prevent Cyclin B1-induced maturation. These results indicate that PKA represses the formation of the starting Cdk1 activity by acting on multiple pathways, some of which being independent of Arpp19 phosphorylation. These discoveries also open new venues of research, aimed at identifying the “early event” controlled by Arpp19. It was proposed that Arpp19 controls the G2/M transition reproduced in interphasic extracts thanks to a crosstalk between the phosphorylations at S109 by PKA and at S67 by Gwl [54]. However, this interplay between phosphorylation sites has not been investigated in prophase oocytes, a time in which Gwl is not active [55]. Further research is therefore required to identify the critical effectors of PKA in the oocytes and characterize their role in the control of the “early events” required for Cdk1 activation.

Our work strengthens a model of Cdk1 activation dependent on a single trigger molecule, progesterone, and a single enzyme, PKA, that is inactivated in response to the hormone. Beyond this bottleneck, however, the dephosphorylation of distinct PKA effectors activate different molecular pathways that, in parallel to each other, converge to activate the starting Cdk1. The unexpected complexity of this network, rather than a linear chain of events, explains why, despite the advances in the understanding of how progesterone causes oocyte maturation in *Xenopus*, nobody succeeded yet in writing down a fully coherent molecular story on the PKA-dependent activation of Cdk1.

## MATERIAL AND METHODS

### Animals

*Xenopus laevis* adult females (Centre de Ressources Biologiques Xénopes, CNRS, France) were bred and maintained according to current French guidelines in the IBPS aquatic animal facility, with authorization: Animal Facility Agreement: #A75-05-25. All *Xenopus* experiments were subject to ethical review and approved by the French Ministry of Higher Education and Research (reference APAFIS#14127- 2018031614373133v2).

### *Xenopus* oocyte handling

*Xenopus laevis* Stage VI oocytes [56] were obtained from unprimed female. *Xenopus* females were anesthetized for 30 minutes in a bath in supplied with 1 g/l MS222 (Sigma) and bicarbonate. Ovarian lobes were collected incubated for 3 h in buffer M (10 mM HEPES pH 7.8, 88 mM NaCl, 1 mM KCl, 0.33 mM Ca(NO3)2, 0.41 mM CaCl2, 0.82 mM MgSO4) in the presence of dispase (0.4 mg/ml) then for 1 h in the presence of collagenase (0.4 mg/ml). After washing with 2 liters of buffer M to eliminate collagenase, the Stage VI oocytes were sorted by size and maintained in M Buffer at 16°C during the experiments. Meiotic maturation was induced by 1 µM progesterone. IBMX was used at 1mM in the external medium and pre-incubated for 1 hr before further manipulation. Oocytes were referred to as NEBD when the first pigment rearrangement was detected at the animal pole.

### Plasmids, mRNA preparation, and recombinant protein purification

The following plasmids were used in this paper: GST-XenCip1 (NM_001094464.1), 6xHis-V5-PKIg, v5-reporter-5’-3’UTRs-Cyclin B1, v5-reporter-5’-3’UTRs-Mos, MBP-GST-flag-HA-V5, GST-Xen-Arpp19-S109D, GST-Xen-Arpp19-S109A, 6xHis-V5-Arpp19-S109D, 6xHis-V5-Arpp19-S109A, and CyclinB1-flag-HA-V5. Cloning was performed using overlap extension PCR or Gibson assembly and Sanger-sequenced. mRNA of all the reporters was *in vitro* transcribed with the mMESSAGE mMACHINE T3 Transcription Kit (Ambion, AM1348); when specified, polyadenylation (150-200 nt) was achieved using the Poly(A) Tailing Kit (Ambion, AM1350). All messages were purified using the MEGAclear Kit (Ambion, AM1908). mRNA concentrations were measured by NanoDrop and message integrity and *in vitro* polyadenylation was evaluated by electrophoresis. The following quantity of mRNAs were injected in each oocyte: v5-reporter-5’-3’UTRs-Cyclin B1 and v5-reporter-5’-3’UTRs-Mos 0.5 ng; MBP-GST-flag-HA-V5 (*in vitro* polyadenylated) 0.1 ng, CyclinB1-flag-HA-V5 (*in vitro* polyadenylated) 15 ng. The recombinant proteins were produced and purified on glutathione-agarose column (G4510, Sigma) for GST-tagged protein or on TALON-Cobalt beads column (635507, Takara Bio) for 6XHIS-tagged protein as previously described [19,26]. The following quantity of recombinant proteins were injected in each oocyte: GST-XenCip1-ORF 60 ng; 6xHis-V5-PKIg 125 ng; GST-Xen-Arpp19-S109D 100 ng, GST-Xen-Arpp19-S109A 100 ng, 6xHis-V5-Arpp19-S109D 100 ng, and 6xHis-V5-Arpp19-S109A 100 ng.

### Oocytes lysates, SDS-page and western blots, antibodies

Oocytes were lysed in 1 oocyte in 10 μL of Extraction buffer (EB: 80 mM β-glycerophosphate pH 7.3, 20 mM EGTA, 15 mM MgCl2). GST-PP1 kinase assay were performed as previously described [26]. SDS-page was performed using 10% or 12% Laemmli gels and proteins were transferred on supported nitrocellulose membranes (Amersham, 10600015) using a semi-dry apparatus as previously described [28]. The following primary antibody were used in PBS+3% BSA+ 0.05% NaN3: Gwl (1:1000, rabbit [19]), Karyopherin (1:2000, goat, Santa Cruz sc-1863), CPEB1 (1:2000, mouse [57]), Cdc6 (1/200 retro-eluted, rabbit [26]), Cyclin B1 (1:200, rabbit [14]), Cyclin B2 (1:1000, mouse, Abcam ab18250), Mos (1:500, rabbit, Santa Cruz Biotechnology Sc-86), pMAPK (1:2000, mouse, Cell Signaling 9106), Erk1/2 (1:2000 each, rabbit, Santa Cruz Biotechnology C-16 and C-14), pT320-pp1 (1:1000 for the detection of the endogenous protein, 1:30000 for the kinase assay, rabbit, Abcam ab62334), pY15-Cdk1 (1:1000, rabbit, Cell Signaling 9111), Cdk1 (1:5000, mouse, Invitrogen MA-91598), Puromycin (1:1000, mouse, Millipore MABE343), and Anti-V5 (1:1000, mouse, Invitrogen 46-0705). The appropriate horse-radish peroxidase (HRP)-labeled secondary antibodies were used at 1:10000 dilution (Jackson Immunoresearch).

### SUnSET

Oocytes were micro-injected with 250 ng of puromycin (Sigma) and incubated 10 min before collection. When oocytes microinjection was performed in MII, the external medium was supplied with 10 mM of EGTA to prevent oocytes activation. Oocytes extracts were analyzed by SDS-page and western blot using the puromycin antibody. Signals were quantified with Image J. Puromycin signals were normalized on the signal from Ponceau Red staining (normalized ratios). The normalized ratios were standardized on the average normalized ratio of each technical replicate. The plot and the statistical analysis were performed with prism.

### Translation reporters

Oocytes were injected with a mixture of 0.5 ng of V5-GST reporter and 0.1 ng of V5-MBP-GST loading control. Oocytes extracts were analyzed by SDS-page and western blot using the V5 antibody. Signals were quantified with Image J. V5-GST signals were first normalized by the V5-MBP-GST one (normalized Ratios).

The normalized ratios were plotted against the time of oocytes collection and a linear regression was performed to calculate the slope of the curve using Excel. The slope represents the rate of translation. The plot and the statistical analysis were performed with prism.

### Size-exclusion chromatography

50 oocytes were homogenized with 450 μL of EB buffer. Extracts were clarified with a 15000 g centrifugation for 20 min and clear supernatants were ultracentrifuged with a TL100 rotor at 53000rpm for 15 min at 4 C. 250 μL of clean extract was injected in gel filtration column (Superose 6), that was previously pre-equilibrated in EB buffer. After 6 ml of dead volume, 500 μL fractions were collected until 30 ml were eluted. Fractions were precipitated by TCA following this protocol. Deoxycholate was added at 0.02% followed by TCA at 10%. Samples were incubated for 1 hr in ice and then centrifuged at 15000g for 20 min at 4 C. Supernatants were removed using a vacuum pump without disturbing the pellets. Pellets were washed with 200 μL of ice-cold acetone and then air dried. Pellets were resuspended in 30 μL of EB 1x and analyzed by SDS-page and western blot.

## ACKNOWLEDGMENTS

We thank all members of the team “Oocyte Biology” for helpful discussions. This work was supported by the National Center for Scientific Research (CNRS, state subsidy), Sorbonne University (state subsidy and “Tremplin - 2022 de la Faculté des Sciences et Ingénierie” to EMD), the National Research Agency (ANR grant 18-CE13-0013-01), and Fondation ARC pour la recherche sur le cancer (ARCPDF22019120001043 to EMD). M.S. received a PhD grant from Sorbonne University (Doctoral School 515, CdV). M.M. thanks Sorbonne University for granting her a CRCT (leave for research or thematic conversions).

