## Supplemental figure 1 for "Regulation of Oocyte Meiotic Maturation: Unraveling the Interplay between PKA Inhibition and Cdk1 Activation"

### Supplementary Figure

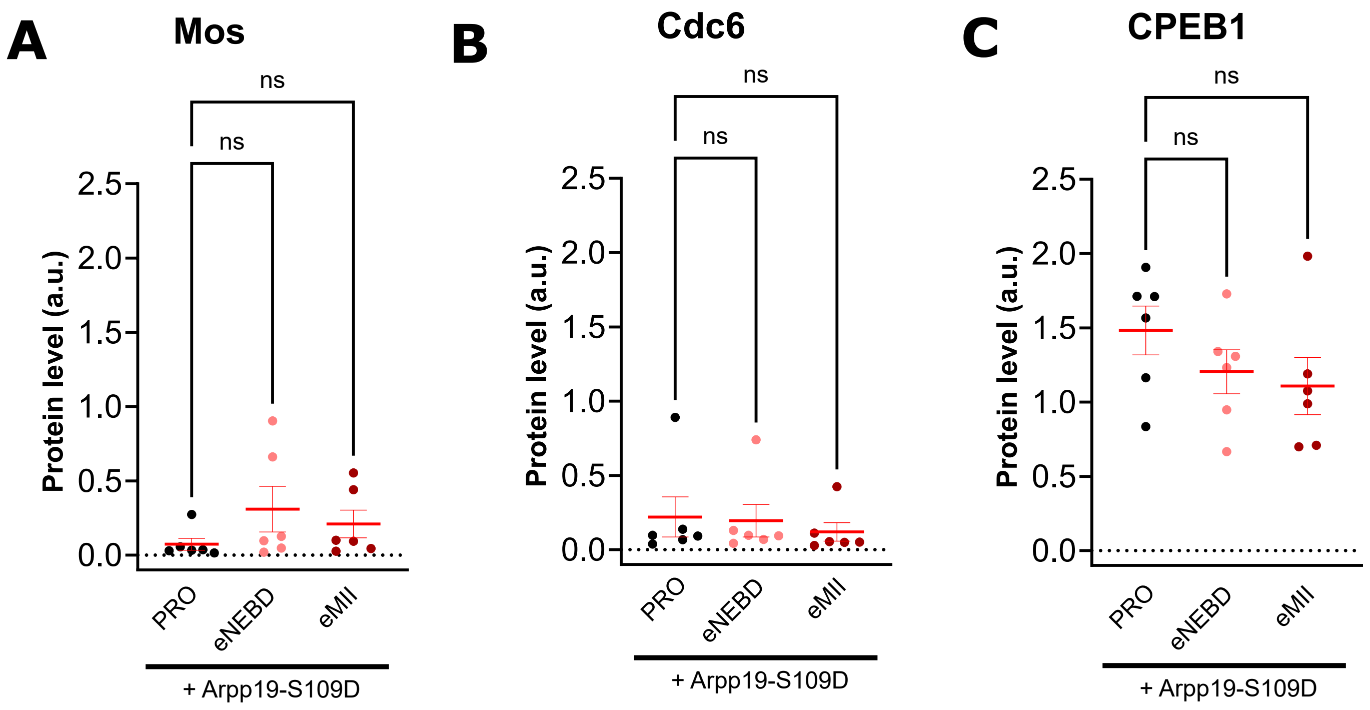


Oocytes were injected or not with either Cip1 or Arpp19-S109D. After overnight incubation, progesterone (Pg) was added in the external medium. Oocytes were collected at prophase (PRO), NEBD, metaphase II (MII) or the equivalent times (eNEBD, eMII) for the Cip1- and Arpp19-S109D injected oocytes. Mos (**A**), Cdc6 (**B**) and CPEB1 (**C**) signals were quantified from oocytes injected with Arpp19-S109D and collected at different times (PRO, eNEBD, eMII). 6 biological replicates were quantified. One-way ANOVAs test with multiple comparisons was performed to evaluate the statistical significance. ns: not significant.
